# The relative importance of biotic and abiotic determinants of temporal occupancy for avian species in North America

**DOI:** 10.1101/555227

**Authors:** Sara Snell Taylor, James Umbanhowar, Allen H. Hurlbert

## Abstract

**Aim:** We examined the relative importance of competitor abundance and environmental variables in determining the species distributions of 175 bird species across North America. Unlike previous studies, which tend to model distributions in terms of presence and absence, we take advantage of a geographically extensive dataset of community time series to model the temporal occupancy of species at sites throughout their expected range.

**Location:** North America.

**Time period:** 2001-2015.

**Major taxa studied:** 175 bird species.

**Methods:** We calculated variation in temporal occupancy across geographic range and used variance partitioning and Bayesian hierarchical models to evaluate the relative importance of 1) the abundance of potential competitors and 2) the environment (elevation, temperature, precipitation, vegetation index) for determining temporal occupancy. We also created a null model to test whether designated competitor species predicted variation in occupancy better than non-competitor species.

**Results:** On average, the environment explained more variance in occupancy than competitor abundance, but this varied by species. For certain species, competitor abundance explained more variance than the environment. Species with larger range sizes, larger range overlap with competitors, and that occurred at higher mean temperatures had a higher proportion of variance explained by the environment than competitor abundance. The abundance of competitor species had a stronger effect on focal species occupancy than non-competitor species in the null model.

**Main conclusions:** Temporal occupancy represents a new way of describing species distributions that is complementary to presence/absence or abundance. Geographic variation in temporal occupancy was explained by both biotic and abiotic drivers, and abiotic drivers explained more variation in temporal occupancy than abundance on average. Species traits also play a role in determining whether variation in temporal occupancy is best explained by biotic or abiotic drivers. The results of our study can improve species distribution models, particularly by accounting for competitive interactions.

## Introduction

Understanding what factors limit species distributions is critical for determining why species persist in their range and what habitat conditions they require (Andrewartha & Birch, 1954; Brown, 1984; Case, Holt, McPeek, & Keitt, 2005; Franklin, 2010; Guisan & Thuiller, 2005). Traditionally, species' distributions have been modelled in terms of presence/absence (Elith et al., 2006; Ferrier, Drielsma, Manion, & Watson, 2002; Phillips, Anderson, & Schapire, 2006) or as a snapshot in time of spatial abundance patterns (Bahn & McGill, 2007; Mönkkönen, Devictor, Forsman, Lehikoinen, & Elo, 2017). An alternative measure of occurrence is temporal occupancy, or the number of years a species was observed at each sampling site over time (Belmaker, 2009; Coyle, Hurlbert, & White, 2013; Taylor, Evans, White, & Hurlbert, 2018). For many taxa, temporal occupancy is strongly bimodal across species at a given site, implying that species either persist in viable or core populations at a site with high temporal occupancy, or are observed only occasionally as transient individuals at low temporal occupancy (Coyle et al., 2013; Taylor et al., 2018; Umaña, Zhang, Cao, Lin, & Swenson, 2017). While temporal occupancy is positively correlated with average abundance in birds, many species maintain persistent populations at low density making this correlation imperfect (Coyle et al., 2013). For this reason, temporal occupancy is a complementary metric to traditional measures for describing species distributions that focuses on geographic variation in population persistence.

Generally, species require specific environmental conditions to succeed in a particular habitat, but often they do not occur everywhere the environment is suitable (Hutchinson 1957; Chesson 2000*a*, Gaston 2003). This distinction between the fundamental and realized niche is usually ascribed to interspecific interactions, such as where a species is outcompeted in parts of its suitable range by a superior competitor (Arif, Adams, & Wicknick, 2007; Connell, 1961; Cunningham, Rissler, Buckley, & Urban, 2016). Recent studies have demonstrated that including interspecific interactions spanning space and time can lead to more complete and accurate species distribution models (Belmaker et al., 2015; Bruno, Stachowicz, & Bertness, 2003; Gotelli, Graves, & Rahbek, 2010; Guisan & Thuiller, 2005) and that interspecific interactions can influence species ranges, even at large scales (Belmaker et al., 2015; Mönkkönen et al., 2017; Ricklefs, 2012). Interspecific competition for desirable habitat and resources may be the most relevant biotic interaction at large temporal and spatial scales of study and is the most studied biotic interaction for shaping species ranges (Sexton, McIntyre, Angert, & Rice, 2009). While many have argued for considering both environmental conditions and biotic factors in order to fully explain species’ distributions, there is not yet consensus on the relative importance of these two categories of drivers, or on the types of species traits that might influence that relative importance.

Here, we seek to quantify the relative importance of biotic and abiotic drivers of temporal occupancy throughout the ranges of North American birds. Because temporal occupancy provides insight into the temporal persistence of populations that a snapshot of abundance cannot, we expect biotic and abiotic predictors to explain more variance in temporal occupancy than abundance. We also examine whether species migratory and foraging traits can help explain why temporal occupancy is better predicted by abiotic variables for some species and biotic interactions for others.

## Methods

### Bird Data

Birds are particularly suitable for modeling species ranges since they are well-studied and there is ample data on their presence over time at large spatial extents (Bennett, Clarke, Thomson, & Mac Nally, 2015; Engler et al., 2017; Palacio & Girini, 2018). We used the North American Breeding Bird Survey (BBS) to characterize geographic variation in species presence and abundance. BBS surveys monitor breeding birds across the continent via a series of fifty 3-minute point counts spaced at 0.8 km intervals along a roadside route (USGS Patuxent Wildlife Research Center 2012). Each survey is conducted during the breeding season (typically June) by a single observer who records all avian species seen or heard along the route. We used the 953 BBS survey routes that were continuously surveyed from 2001-2015, and we excluded species that were poorly sampled by the survey design such as waterbirds, raptors, and nocturnal species (Robbins, Bystrak, & Geissler, 1986). We identified 175 focal land bird species for analysis based on the following criteria: 1) species were present on at least 50 BBS routes over the 15 sampling years, and 2) species were observed at more than 30 percent of the survey sites within their geographic ranges (based on BirdLife International shapefiles, www.birdlife.org) over a ten-year period (Hurlbert & White, 2007).

### Biotic Drivers

For each focal species, competitor species were identified as those species in the same family with any area of overlapping geographic range and similar body size (within two-fold of the body mass of the focal species). A comprehensive list of focal and associated competitor species is included in Table S1. In some cases, potential competitors from outside the focal species’ family were included when such interactions were specifically described in natural history accounts (e.g. American Redstart, *Setophaga ruticilla* and Least Flycatcher, *Empidonax minimus*, Sherry 1979). From the list of competitors for each focal species, a main competitor was identified as the species with the greatest percentage of breeding range overlap with the focal species based on BirdLife International shapefiles.

At each survey route *r*, we calculated two indices of competitive pressure, *c*_*main*_ and *c*_*sum*_, based on the proportional abundance of competitors relative to the focal species:

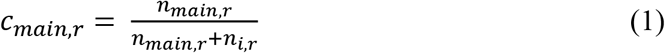

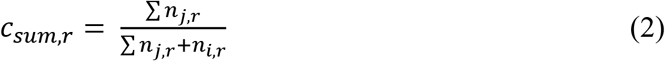

where *n*_*i*,*r*_ and *n*_*main*,*r*_ are the abundances of focal species *i* and the main competitor of species *i*, respectively, on route *r*, and *n*_*j*_ is the abundance of the *j*th competitor of species *i*. We use these scaled indices of competitor abundance rather than raw abundances to account for variation in absolute densities over the broad geographic gradients examined.

### Abiotic Drivers

Long term normals for mean annual temperature and mean annual precipitation were acquired from WorldClim (Hijmans, Cameron, Parra, Jones, & Jarvis, 2005) at 2.5-minute resolution, and a 30-m digital elevation model was acquired from the National Elevation Dataset (NED from USGS). Normalized difference vegetation index (NDVI) was acquired from the Global Inventory Mapping and Monitoring Studies database (GIMMS). The NDVI data was 250 meter, monthly resolution, averaged over the summer months of June, July, and August (https://gimms.gsfc.nasa.gov/MODIS/).

For each BBS route, the mean of each environmental variable was calculated within a 40 km buffer to ensure that the entire 40 km route was encompassed. For each species *i*, the environmental centroid or species’ optimum, 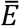, for each environmental variable *k* was calculated by weighting the environmental value at each BBS route by the focal species’ average abundance at that site (*n*_*ir*_):

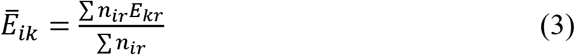

Environmental values for each variable *k* were then transformed into *z*-scores describing the absolute deviation, *D*, from the optimum for species *i* at route *r* as follows:

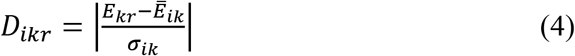

where *σ*_*ik*_ is the abundance-weighted standard deviation of environmental values across all routes at which species *i* is present. We took the absolute value of these environmental deviations because we are not concerned with the direction of deviations, simply their magnitude. Environmental deviation values were calculated for temperature, precipitation, elevation, and NDVI across the set of BBS routes at which each focal species was expected to occur based on range map shapefiles. Higher values of this deviation metric indicate a greater deviation from the species’ environmental optimum.

### Analysis

Temporal occupancy was calculated at each BBS survey route for each focal species as the fraction of sample years between 2001-2015 during which the species was observed. Each focal species was also classified with respect to migratory class, family, and diet category based on Hurlbert and White (2007). Breeding range centroids were calculated based on the BirdLife International shapefiles using the R package “Gtools” (Warnes, Bolker, & Lumley, 2015).

For each focal species, we used variance partitioning to quantify the amount of variance in temporal occupancy that could be uniquely explained by either the set of environmental variables or scaled competitor abundance (*c*_*main*_ or *c*_*sum*_), as well as the shared variance component that could not be uniquely ascribed to either variable class. We also developed a hierarchical Bayesian model to understand how competitor abundance and environmental deviation from the optimum influenced focal species occupancy. The model used focal occupancy as the response variable and scaled competitor abundance and deviation measures for each of the four abiotic variables, allowing for random slopes and intercepts by species:

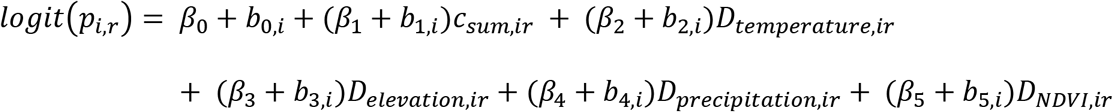

where p_*ir*_ represents the probability of a given focal species *i* appearing any given year at site *r*, *β*_*j*_ are the population level effect coefficients, and *b*_*j*,*i*_ are the species level random effects. We assumed that the random effects were distributed *b*_*i*_ ~ Mulivariate normal(0,∑), where ∑ is an unstructured variance covariance matrix. Observed occupancies we assumed were distributed *T*_*ir*_ ~ Binomial(*p*_*ir*_,*n*_*r*_), where *n*_*r*_ were the number of years sampled at site *r*. The model was estimated using the brms package using the default priors in version 2.7.0 (Bürkner, 2017).

We quantified the extent to which temporal occupancy for a given species was better predicted by biotic or abiotic variables using the ratio, *R*_*C*_, of the competition variance component to the sum of the competition and abiotic environment variance components. Values of *R*_*C*_ above 0.5 indicate that of the variance in temporal occupancy explained, more variance is explained by competitor abundance than environment, while values below 0.5 indicate that more variance is explained by the environment than competitor abundance. We explored whether several species traits could explain interspecific variation in *R*_*C*_ using logit transformed linear models. We examined how *R*_*C*_ varied with migratory and trophic group categories, weighting each species in the analysis based on the total number of BBS routes on which it occurred over the 15-year period. The rationale for weighting species in this way is that species with larger ranges and hence larger spatial sample sizes should have better R^2^ estimates. We also evaluated a separate model of continuous predictors of *R*_*C*_ including each species’ environmental centroids for temperature, precipitation, elevation, NDVI, range size, and proportional area of overlap with competitors. Proportional area of overlap was calculated by summing the total area each focal species overlapped with a potential competitor species, divided by the focal range area. Because range overlap is summed across multiple competitor species, it may exceed one. A list of focal species and the traits used in these analyses is provided in Table S2.

### Competitor null model

To better interpret the amount of variance in temporal occupancy explained by competitor abundance, we conducted a null model analysis examining the variance explained by the scaled abundance of non-competitor species. For our purposes, non-competitor species included the subset of all species in our dataset from a different family as the focal species (with some exceptions as noted), or from the same family but where body size differed from the focal species by 2-fold or more, and with some overlap in geographic range. For each focal species where main competitor abundance had a strong effect (R^2^ ≥ 10%, *n* = 61), we conducted separate linear regressions predicting focal species occupancy based on the scaled abundance of each non-competitor (based on Eq. 1). The number of non-competitor species evaluated for each focal species varied between 116 and 274. Any variance explained by non-competitor species presumably reflects indirect habitat associations rather than competitive effects, and thus provides a benchmark for interpreting the variance explained and the effect size of the competitors on the focal species. We expected a stronger negative relationship between focal occupancy and abundance for the main competitor than for non-competitors, and we calculated the proportion of non-competitor species with a higher R^2^, or more negative parameter estimate, than observed for the main competitor.

## Results

Species differed substantially in the total amount of variance in temporal occupancy explained and in whether competitor abundance or environmental factors explained more variance (Figure 1). The median total variance explained by both sets of predictors together was 31% across all 175 species, and abiotic factors uniquely explained more variance in temporal occupancy than summed competitor abundance on average (12% compared to 8%; Figure 1a, Table S3). The ratio *R*_*C*_, which describes the relative amount of variance explained by competitor abundance, spanned a wide range of values. While a few species had high values, the bulk of the distribution fell under 0.5 (median = 34%) indicating stronger predictive power of abiotic over biotic variables. Using the abundance of a single main competitor rather than the summed abundance of all competitors decreased the median variance explained by competitor abundance (3%, compared to 8% for all competitors), while the environmental median variance component increased (16%; Figure 1b, Table S4). Total variance and the ratio *R*_*C*_ decreased (28% and 18%, respectively). Unless otherwise specified, subsequent analyses refer to the summed abundance of all competitors to characterize competitive pressure.

**Figure 1.**
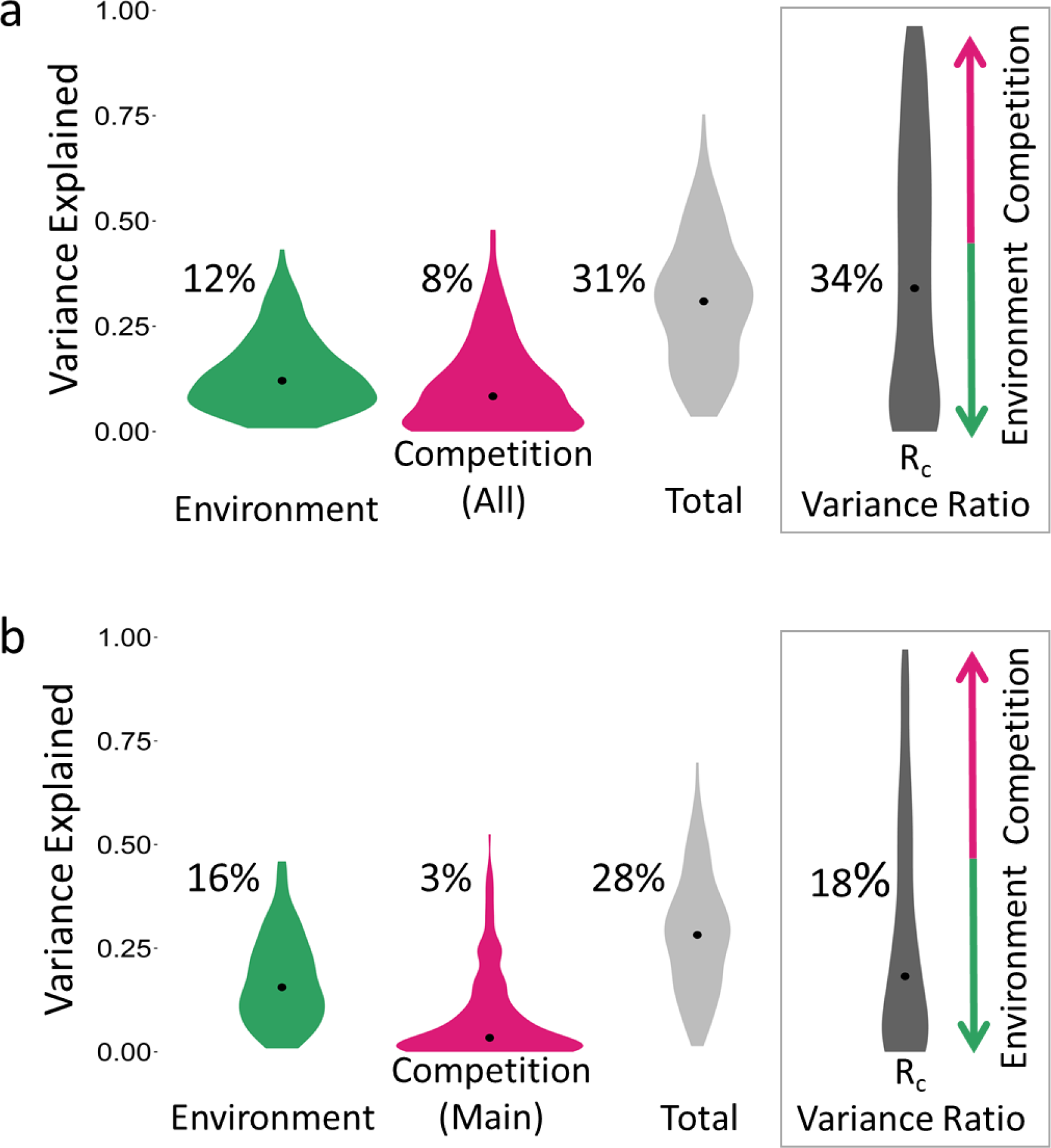
(a) Violin plots demonstrating the distribution of unique variance in focal occupancy explained by environmental variables, the unique variance explained by the summed abundance of all competitors,, the total variance explained by a model with both sets of variables, and R_c_, the competition variance component divided by the sum of the competition and environment variance components. Median values are noted within each distribution. (b) Violin plots demonstrating the distribution of unique variance in focal occupancy explained by environmental variables, the unique variance explained by the abundance of the main competitor, the total variance explained by a model with both sets of variables, and the variance ratio, R_c_.

The variance partitioning results for the 15 species that had the most variance in temporal occupancy explained by competitor abundance and by the environment, respectively are shown in Figures 2 and 3 (full variance partitioning results and model output for all species are provided in Tables S3 – S6). The range of unique variance explained for these top species was similar for competitor abundance (28-48%, Figure 2) and environment (29-43%, Figure 3) variance components, and species that were well explained by one variable category typically had little variance explained by the other. Shared variance that could not be uniquely ascribed to either competitors or the environment was higher for those species best explained by competitor abundance. Even though the median variance component explained by the environment was higher than the median variance component explained by competitors, the maximum variance component explained by competitors was higher than the maximum variance component explained by the environment. Phylogeny was unrelated to the amount of explained variance, as nine different families were represented within the top 15 species ranked by environment-explained variance and 11 families were represented in the top 15 species ranked by competitor-explained variance.

**Figure 2.**
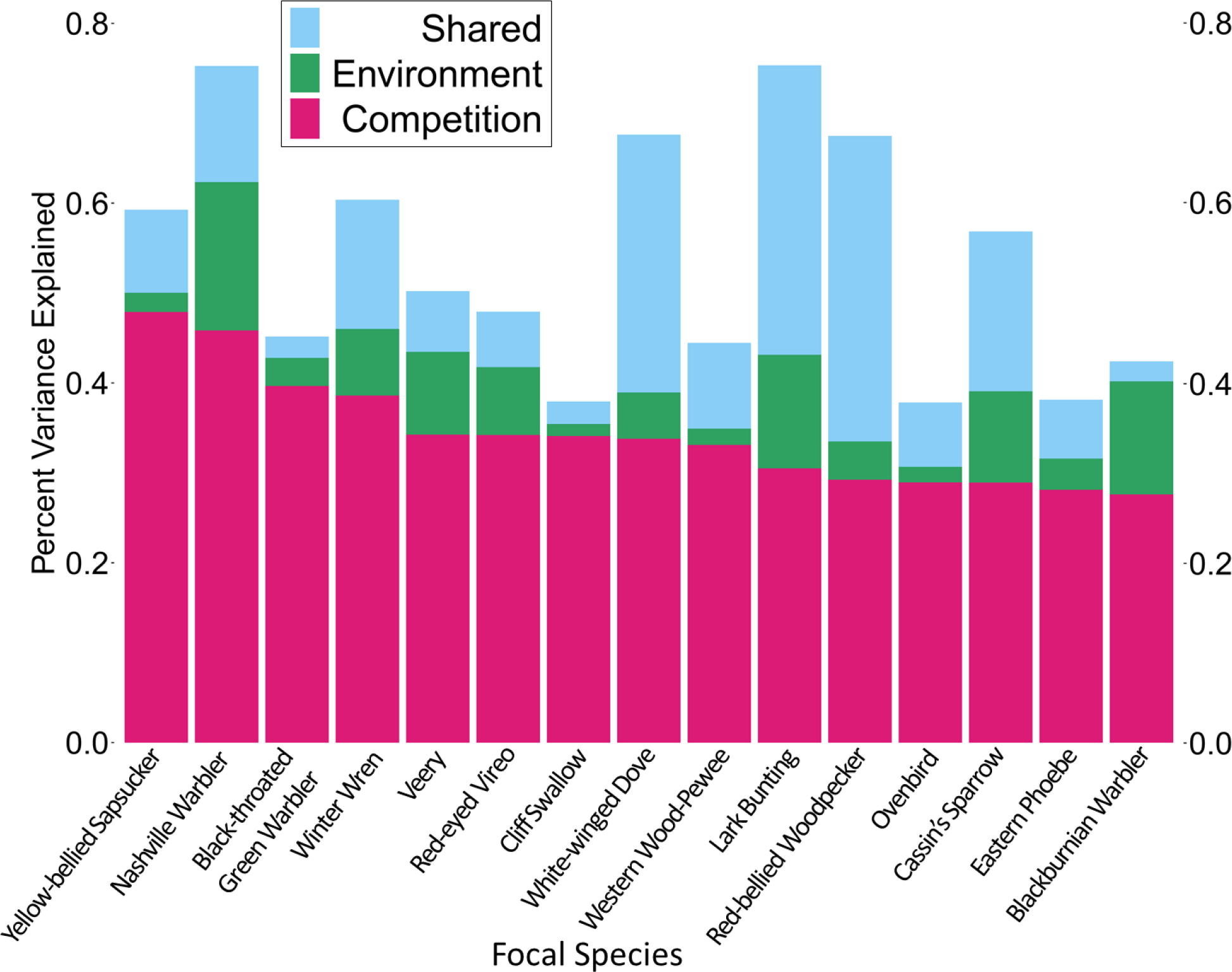
The top 15 focal species ranked by variance explained by scaled competitor abundance. Variance partitioning results illustrating the variance uniquely explained by competition (pink), uniquely explained by the environment (green), and the shared variance component that cannot be uniquely ascribed to either class of variables (blue).

**Figure 3.**
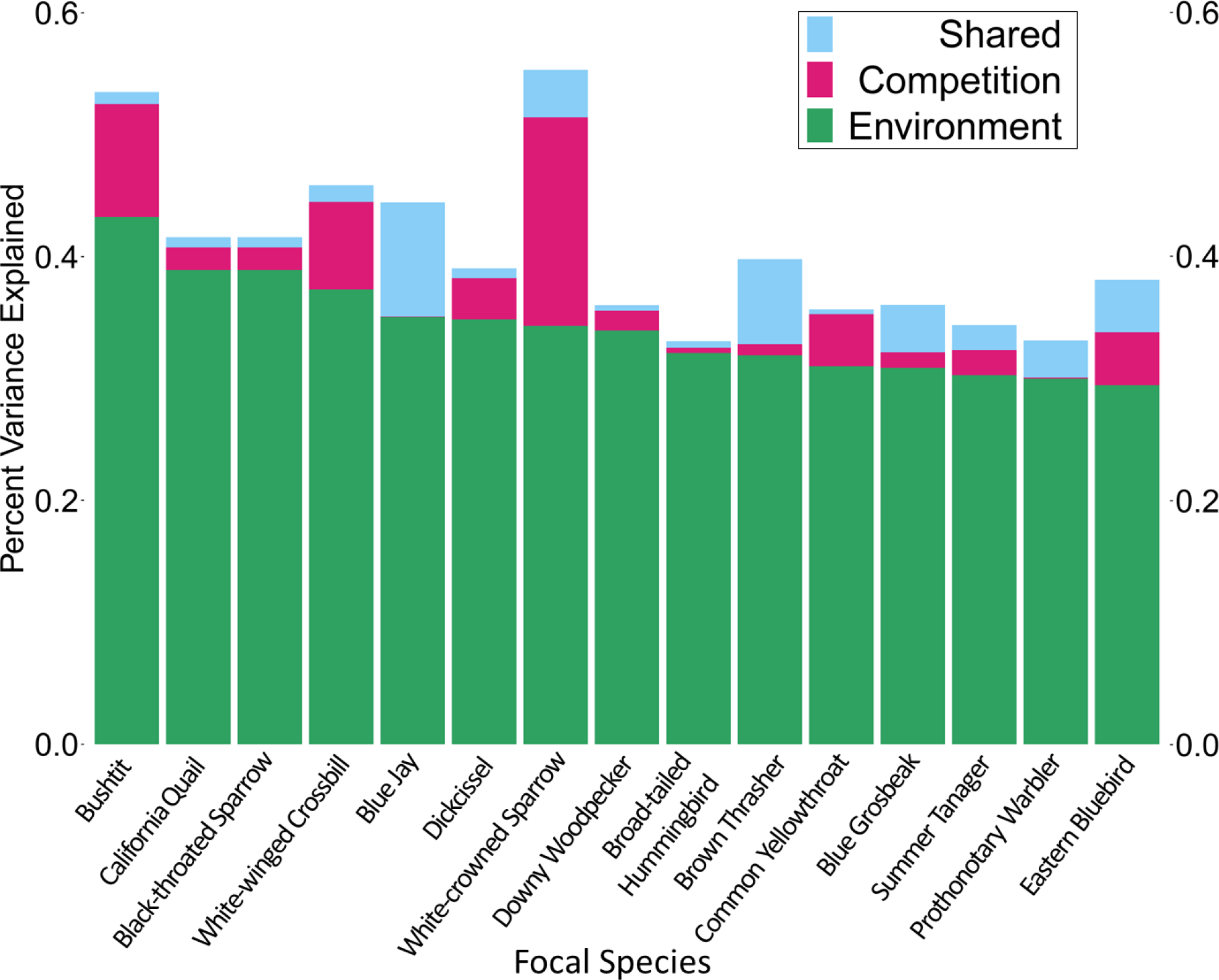
The top 15 focal species ranked by variance explained by deviation from optimal environmental conditions. Variance partitioning results illustrating the variance uniquely explained by uniquely explained by the environment (green), competition (pink), and the shared variance component that cannot be uniquely ascribed to either class of variables (blue).

Focal species occupancy was lower at sites that deviated farther from the species’ environmental optimum for all environmental variables, and occupancy decreased as relative competitor abundance increased (Table 1). Across all variables, there was wide variation in the relationship between the environment, competitor abundance, and focal species occupancy (Table 1). Temperature (slope = −0.45 [−0.53, −0.37]) and NDVI (slope = −0.27 [−0.32, −0.21]) had stronger negative relationships with focal occupancy than elevation (slope = −0.15 [−0.23, −0.07]) or precipitation (slope = −0.16 [−0.23, −0.09]; Table 1). Competitor abundance (slope = −2.00 [− 2.56, −1.53]) had a strong negative relationship with focal occupancy, although its slope is not directly comparable to the slopes of environmental deviation since competitor abundance is scaled from zero to one. All 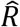 were between 1.00 and 1.03. Full model output is available as supplementary material (Table S7 for all competitors, Table S8 for main competitors).

**Table 1.** Bayesian hierarchical model results for the fixed effects of the absolute value of the environmental z-score (deviation from the environmental centroid) and competitor abundance on focal species occupancy. Species was included as a random effect allowing for random slopes and intercepts.

| | Mean | Credible Interval | | $\sigma$ | Credible Interval of $\sigma$ | |
| --- | --- | --- | --- | --- | --- | --- |
|  |  | Lower 95% | Upper 95% |  | Lower 95% | Upper 95% |
| Intercept | 3.26 | 2.75 | 3.88 | 3.74 | 3.35 | 4.18 |
| Competitor |  |  |  |  |  |  |
| Abundance | -2.00 | -2.56 | -1.53 | 3.38 | 3.02 | 3.78 |
| Temperature | -0.45 | -0.53 | -0.37 | 0.50 | 0.45 | 0.56 |
| NDVI | -0.27 | -0.32 | -0.21 | 0.35 | 0.31 | 0.39 |
| Elevation | -0.15 | -0.23 | -0.07 | 0.52 | 0.47 | 0.58 |
| Precipitation | -0.16 | -0.23 | -0.09 | 0.48 | 0.43 | 0.54 |

We found that abiotic variables collectively explained approximately twice as much variance in temporal occupancy as they did in abundance across sites (Figure 4, green circles). In contrast, abundance was better predicted by competitor abundance on average, with the majority of points falling above the 1:1 line (Figure 4, pink triangles). Note that for main competitors, temporal occupancy and abundance were equally well predicted by competitor abundance (Figure S1). The total variance explained by the combination of biotic and abiotic variables was only slightly greater on average when predicting abundance compared to temporal occupancy, although individual species differed substantially (Figure 4, gray crosses). Total variance was slightly greater when predicting temporal occupancy compared to abundance when using only a main competitor abundance (Figure S1). For example, 75% of the total variance in temporal occupancy could be explained for the Lark Bunting compared to only 52% of the total variance in abundance, while for the Worm-eating Warbler, only 16% of the variance in occupancy could be explained compared to 51% for abundance. Full variance partitioning results and model output for all species predicting focal abundance rather than occupancy is available in Tables S9 – S12.

**Figure 4.**
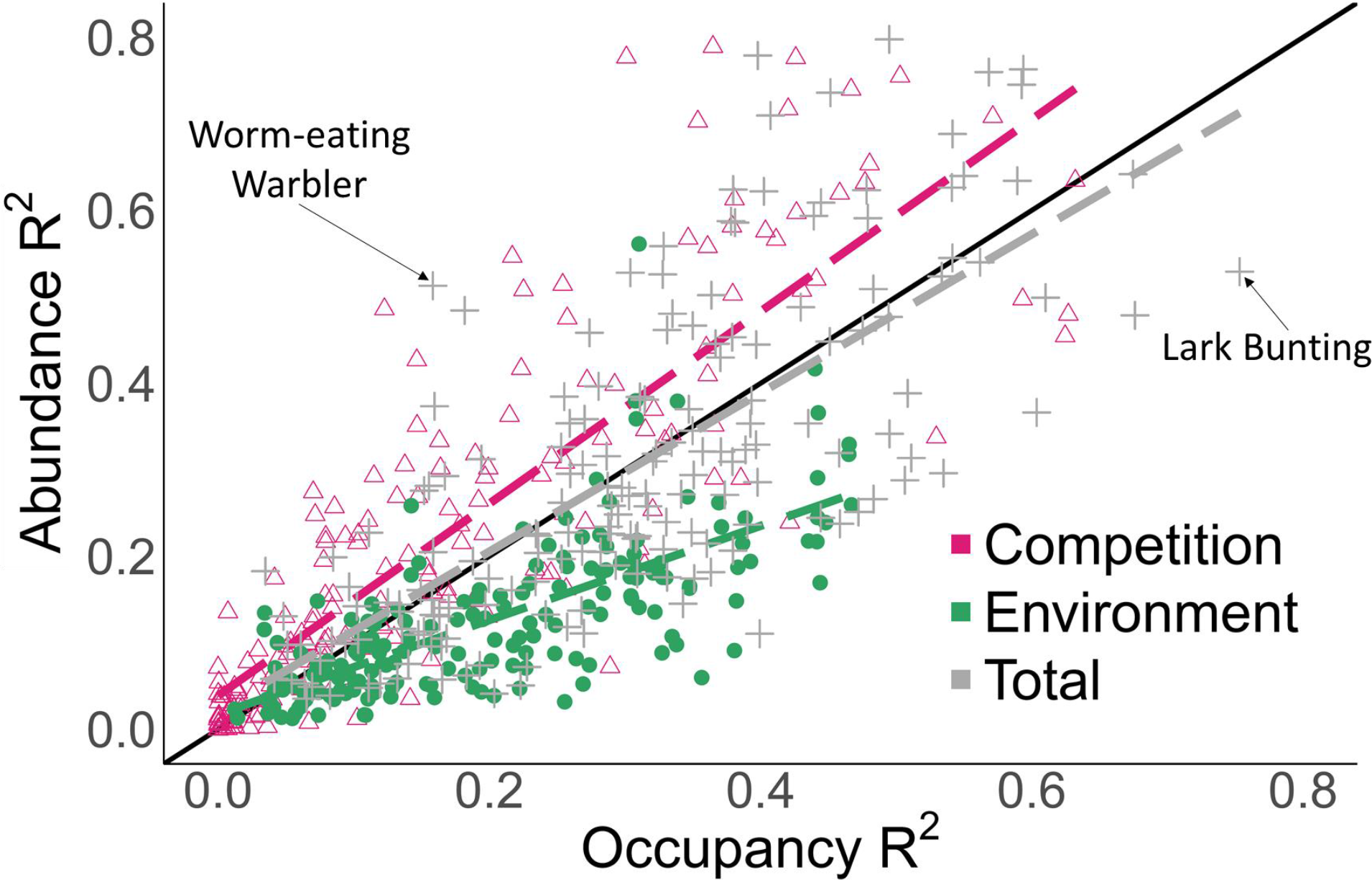
The ability of environmental variables (green circles), competitor abundance (pink triangles), or both combined (grey crosses) to predict spatial variation in temporal occupancy (x-axis) compared to spatial variation in abundance (y-axis) based on linear model R^2^s (and not unique variance components, as portrayed in Figures 1–3). Black line represents the 1:1 line. Dashed lines indicate linear regressions through each of the three sets of predictor variables.

Species differed in the relative explanatory power of biotic or abiotic variables for predicting temporal occupancy, measured by the ratio *R*_*C*_, and we examined whether species traits could explain this variation. On average, more variance in occupancy was explained by the environment than competitor abundance regardless of trophic group (insectivores, insectivore/omnivores, granivores, and omnivores, p > 0.12) or migratory status (residents, short-distance migrants, neotropical migrants, p > 0.10). A model using several continuous traits and predictors explained 17% of the variance in *R*_*C*_. Species with larger ranges and higher mean temperatures had a greater proportion of explained variance due to the environment (i.e., lower *R*_*C*_ values) compared to competitor abundance (Table 2). Additionally, as range overlap between species and their competitors increases, the proportion of explained variance due to competition also increased. Mean elevation, precipitation, and NDVI had no effect on why some species are relatively better predicted by the environment than others (Table 2).

**Table 2.**
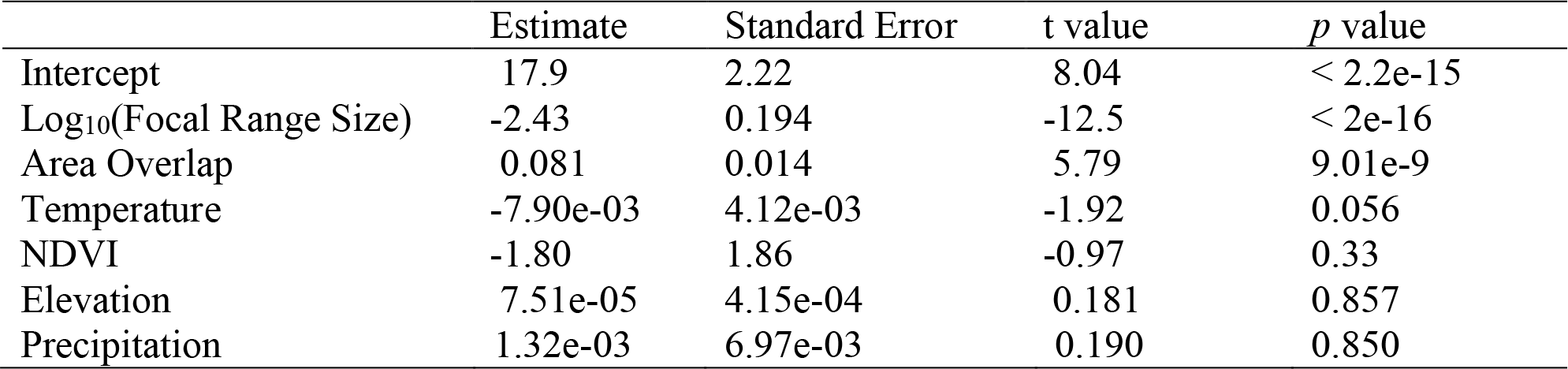
Linear model of the effect of continuous traits (range size, proportional area of overlap between focal species and all competitors, and abundance-weighted average temperature, precipitation, elevation, and NDVI calculated across each species’ range) on logit-transformed *R*_*C*_, the competition variance component divided by the sum of the competition and environment variance components.

In the null model, we modelled the temporal occupancy of each focal species as a function of the scaled abundance (Eq. 1) of each non-competitor species in our dataset. As an example, the Yellow-bellied Sapsucker (*Sphyrapicus varius*) was compared to 154 non-competitors that were able to explain a median of 3% of the variance in occupancy, compared to 57% explained by its assigned main competitor, the Hairy Woodpecker (*Leuconotopicus villosu*; Figure 5a). Additionally, the Hairy Woodpecker had a much stronger negative effect on Yellow-bellied Sapsucker temporal occupancy compared to the median effect size of non-competitors (−12.5 versus −3.2, Figure 5b). For the 61 focal species with a strong (R^2^ ≥ 10%) effect of main competitor abundance, only a small proportion of null non-competitors could explain more variance in temporal occupancy than the main competitor (Figure 5c). Across these focal species, the explained variance of the putative main competitor was much higher than the median variance explained by non-competitors (paired *t*-test, p < 2e-16). Similarly, we found that few non-competitors had effect sizes that were more negative than the effect size of the main competitor (paired *t*-test, p < 2e-16; Figure 5d).

**Figure 5.**
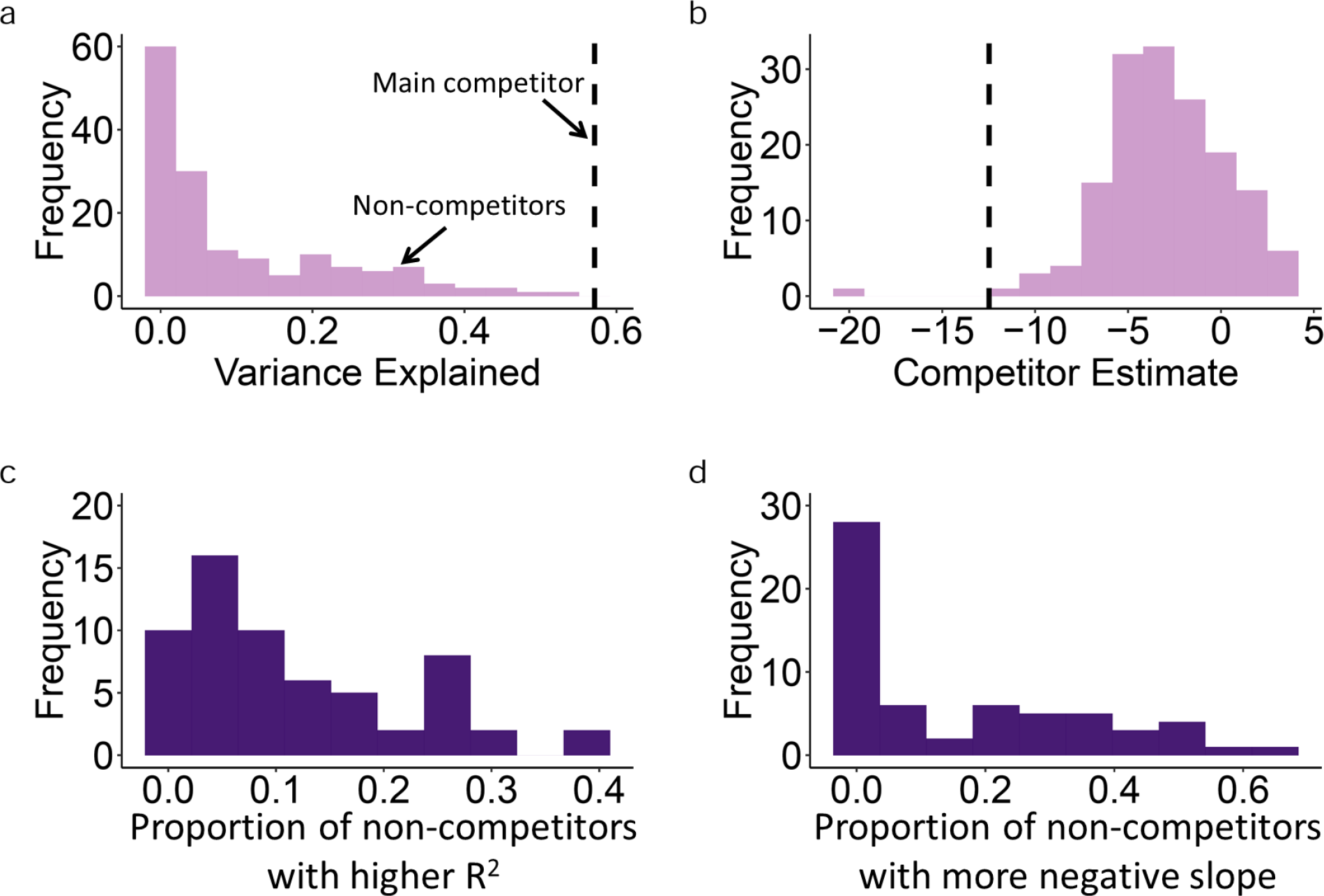
(a) Histogram of 154 non-competitors of example focal species Yellow-bellied Sapsucker R^2^ (median = 0.03); black dashed line is R^2^ of assigned main competitor (Hairy Woodpecker; Competition R^2^ = 0.57). (b) Histogram of 154 non-competitors of Yellow-bellied Sapsucker estimate (median = −3.2); black line is estimate of assigned main competitor (median = −12.5). (c) Histogram of the proportion of non-competitor species with R^2^ values matching or exceeding the main competitor R^2^ when predicting focal species occupancy. (d) Histogram of the proportion of non-competitor species with estimates matching or exceeding the main competitor estimate when predicting focal species occupancy.

## Discussion

Temporal occupancy reflects the persistence of a population over time and varies broadly throughout a species’ range. We found that both biotic and abiotic variables explain a large fraction of the geographic variation in temporal occupancy for any given species. While abiotic environmental variables typically explained more variation than the abundance of interspecific competitors, there were many species for which the opposite was true, as well as some species whose occupancy was poorly explained by all variables considered. For most species, environmental variables could better predict spatial patterns of temporal occupancy than they could spatial patterns of abundance, which have been more traditionally examined.

Abiotic factors have historically received the most attention in the literature for explaining species distributions (Andrewartha & Birch, 1954; Gaston, 2003; Sexton et al., 2009). Here, we found that species tend to have the highest temporal occupancy in environments that are closest to their range-wide environmental centroids, with decreasing occupancy in environments that are most different from the centroid conditions. Overall, temperature had the strongest effect on focal temporal occupancy of the environmental variables considered, but no single variable consistently explained more variance in occupancy compared to the other environmental variables in the single-species models. For example, the Bushtit had the most variance in temporal occupancy explained by the environment (43%), with the strongest effects of temperature, followed by NDVI, and to a lesser extent elevation and precipitation. In contrast, occupancy of the Black-throated Sparrow only exhibited a strong negative relationship with NDVI. These examples demonstrate variation in the exact environmental determinants of occupancy for individual species, but collectively, environmental variables explained more variance in occupancy than competitor abundance on average.

Ecologists increasingly recognize that biotic factors may also be important in shaping distributions over broad geographic scales (Araújo & Rozenfeld, 2014; Belmaker et al., 2015; Bruno et al., 2003; Mönkkönen et al., 2017). Even though abiotic variables generally explained more variance in temporal occupancy for the focal species, the maximum amount of variance that could be explained by competitor abundance (48% for Yellow-bellied Sapsucker) was greater than the maximum amount of variance that could be explained by the environment (43% for Bushtit). The observed decline in temporal occupancy as competitor abundance increased is what we would expect if increasing competition made it more difficult for the focal species to persist at certain sites. We see these effects regardless of whether we used the abundance of a single main competitor species or the summed abundance of all potential competitors.

Additionally, these observed negative effects of competitor species were stronger and explained more variance than those of non-competitors, supporting the interpretation of competition (past or present) rather than associations due simply to differences in habitat preferences.

Nevertheless, there are some limitations to our approach and caveats in interpretation. Field studies quantifying the strength and consequences of interspecific competition in birds are time intensive (Dhondt, 2012) and have not been conducted for most species. To assign potential competitors to 175 focal species in a standardized fashion, we used a simple set of criteria: that they be from the same family (unless there was literature demonstrating a non-familial competitive relationship), that they be similar in body size, and that their geographic ranges overlap. These selected species may include species that do not strongly compete with the focal species, introducing noise and potentially resulting in the low explained variance due to competitor abundance for some species. Using information on foraging behavior as well as morphological traits like bill, wing, and leg dimensions could possibly refine competitor assignments in future studies. Conversely, focal species may compete for resources with heterofamilial species that we did not consider, or even with other taxonomic groups (Brown, Davidson, & Reichman, 1979).

Another limitation is while negative effects of competitor abundance on focal species occupancy are consistent with competitive interactions, they may also be consistent with divergent habitat preferences that lead to negative correlations in space. Such divergent habitat preferences may or may not result from past selection (Connell, 1980). Consider the Yellow Warbler (*Setophaga petechia*), whose broad geographic range leads to high range overlap — and therefore assignment of “main competitor” status — with many other warbler species in our dataset (Table S1). Despite its broad geographic range, the Yellow Warbler preferentially breeds in wet, deciduous thickets and is commonly associated with willows (Lowther, Celada, Klein, Rimmer & Spector, 1999). For other warbler species, a negative correlation with Yellow Warbler abundance may simply reflect negative associations with Yellow Warbler’s preferred habitat rather than evidence for ongoing competition. This is likely the case for most of the warbler species whose occupancies were strongly predicted by Yellow Warbler abundance, given the stated habitat preferences in their respective Birds of North America species accounts (Rodewald 2018). In some cases, the variance explained by abundance of the main competitor may actually reflect finer-scale habitat associations rather than competition. That said, unless a species differs in habitat preference from all other members of its family, the use of the summed abundance of all potential competitors should minimize the influence of this alternative interpretation.

The scale at which we conducted our analyses likely affected observed occupancy patterns and potentially the determinants of those patterns (Jenkins, White, & Hurlbert, 2018; Taylor et al., 2018). Because we used environmental and community data collected at the scale of ~40 km, we can only make inferences related to competition at the landscape scale. Competitive interactions have certainly been documented at these scales and larger (Belmaker et al., 2015; Gotelli et al., 2010), however, our analysis was incapable of detecting the interspecific competition that occurs at much finer scales, as demonstrated in classic studies of local niche partitioning (Dhondt, 2012; MacArthur, 1957; Morse, 1980). As such, finding that competitor abundance explains little variation in temporal occupancy for any particular species clearly does not imply that competition is altogether unimportant for that species.

We examined temporal occupancy as a response that varied across a species’ geographic range in contrast to previous studies that have examined spatial variation in abundance (Araújo & Rozenfeld, 2014; Bahn & McGill, 2007; Brown, 1984; Mehlman, 1997) or presence/absence (Elith et al., 2006; Ferrier et al., 2002; Phillips et al., 2006). Environmental variables were better able to predict spatial variation in temporal occupancy than spatial variation in abundance, while competitor abundance better predicted focal species abundance than focal species occupancy. This difference in explanatory power based on the type of predictor highlights important differences in the ecological information encoded in occupancy versus abundance. Because temporal occupancy integrates how a species interacts with its environment over time, it may produce a more accurate characterization of that species’ fundamental niche. Temporal occupancy may also help distinguish between sites where a species shows up as a rare and infrequent transient species (Taylor et al., 2018) as opposed to a rare but persistent member of the community. This further implies that species distribution models, which traditionally use environmental variables to predict presence or abundance, might have improved performance predicting temporal occupancy. Conversely, summed competitor abundance explained more variance in spatial abundance patterns than spatial occupancy patterns. Given that a species can persist under a given set of environmental conditions, the average population size it is able to obtain there may be in part due to the abundance of other competitors. Thus, temporal occupancy and abundance appear to be complementary measures of species distribution that may each help characterize a species’ realized niche. More work is needed to understand what types of species and in which environmental contexts occupancy will be most influenced by biotic and abiotic factors, and how such an understanding might ultimately inform habitat suitability models for conservation.

## Acknowledgements

We are grateful to M. Jenkins, R. Burger, and G. Di Cecco for comments on this manuscript and study design, and to the volunteers who collect data for the North American Breeding Bird Survey. Thanks to C. Tucker for suggestions regarding the null model for competitor abundance. This work was made possible by funding from the National Science Foundation through grant DEB-1354563 to Allen H. Hurlbert and Ethan P. White and by the Gordon and Betty Moore Foundation’s Data-Driven Discovery Initiative through grant GBMF4563 to Ethan P. White. The funders had no role in study design, data collection and analysis, decision to publish, or preparation of the manuscript.

## Data accessibility statement

The data and code associated with this study will be available on Zenodo and the data are available as a zip file in the Supporting Information.

