## Supplemental Tables 1-12 for "The relative importance of biotic and abiotic determinants of temporal occupancy for avian species in North America": Data S1.docx

List and descriptions of supplemental tables references in manuscript.

### File list (files found within DataS1.zip)

DataS1.doc

TableS1.txt

TableS2.txt

TableS3.txt

TableS4.txt

TableS5.txt

TableS6.txt

TableS7.txt

TableS8.txt

TableS9.txt

TableS10.txt

TableS11.txt

TableS12.txt

**Description**

DataS1.doc – Document describing data included in supplemental tables.

TableS1.txt – Table of all focal species and assigned competitors considered in our analyses. Focal species common name, scientific name, id code (AOU), and family are listed, along with all competitors for each focal species, with the main competitor denoted by a 1.

TableS2.txt – Table of all focal species and traits used in our analyses: focal species family, body mass, migratory class, trophic group, range size of focal species, proportion of overlap in range with competitors, mean optimal temperature, mean optimal precipitation, mean optimal elevation, and mean optimal NDVI.

TableS3.txt – Tabular output of temporal occupancy variance partitioning analysis using all competitors. Each row represents one focal species and the relative contribution of the environmental variables, competitor abundance, shared variance, total variance (environment plus competitor plus shared variance), number of sites (n), and the variance ratio (R_c_) included in the analysis.

TableS4.txt – Tabular output of temporal occupancy variance partitioning analysis using a single main competitors. The linear models used to create the table are the same as for all competitors, however in this table the scaled competitor abundance only contains one assigned main competitor rather than all competitors. Each row represents one focal species and the relative contribution of the environmental variables, competitor abundance, shared variance, total variance (environment plus competitor plus shared variance), number of sites (n), the variance ratio (R_c_), and the main competitor included in the analysis.

TableS5.txt – Tabular output of temporal occupancy variance partitioning analysis for all competitors. Each row contains the estimate, R^2^, and p-value for occupancy and abundance for each focal species.

TableS6.txt – Tabular output of temporal occupancy variance partitioning analysis for a single main competitor. Each row contains the estimate, R^2^, and p-value for occupancy and abundance for each focal species. Note that eight species did not have greater than two occurrences of having expected presence with their main competitor, so their results are NA since we could not perform a linear model on them. These species are included in other parts of the analysis.

TableS7.txt – Tabular output of Bayesian hierarchical model output of random slopes and intercepts grouped by family using all competitors. Each row represents a focal species intercept, environmental factor, and/or scaled competitor output.

TableS8.txt – Tabular output of Bayesian hierarchical model output of random slopes and intercepts grouped by family using a single main competitor. Each row represents a focal species intercept, environmental factor, and/or scaled competitor output.

TableS9.txt – Tabular output of abundance variance partitioning analysis using all competitors. Each row represents one focal species and the relative contribution of the environmental variables, competitor abundance, shared variance, total variance (environment plus competitor plus shared variance), number of sites (n), and the variance ratio (R_c_) included in the analysis.

TableS10.txt – Tabular output of abundance variance partitioning analysis using main competitors. The linear models used to create the table are the same as for all competitors, however in this table the scaled competitor abundance only contains one assigned main competitor rather than all competitors. Each row represents one focal species and the relative contribution of the environmental variables, competitor abundance, shared variance, total variance (environment plus competitor plus shared variance), number of sites (n), the variance ratio (R_c_), and the main competitor included in the analysis.

TableS11.txt – Tabular output of abundance variance partitioning analysis for all competitors. Each row contains the estimate, R^2^, and p-value for occupancy and abundance for each focal species.

TableS12.txt – Tabular output of abundance variance partitioning analysis for main competitors. Each row contains the estimate, R^2^, and p-value for occupancy and abundance for each focal species.
