## Supplementary figures and images for "The relative importance of biotic and abiotic determinants of temporal occupancy for avian species in North America"

### Figure S1 _occvabun_lines_main.pdf

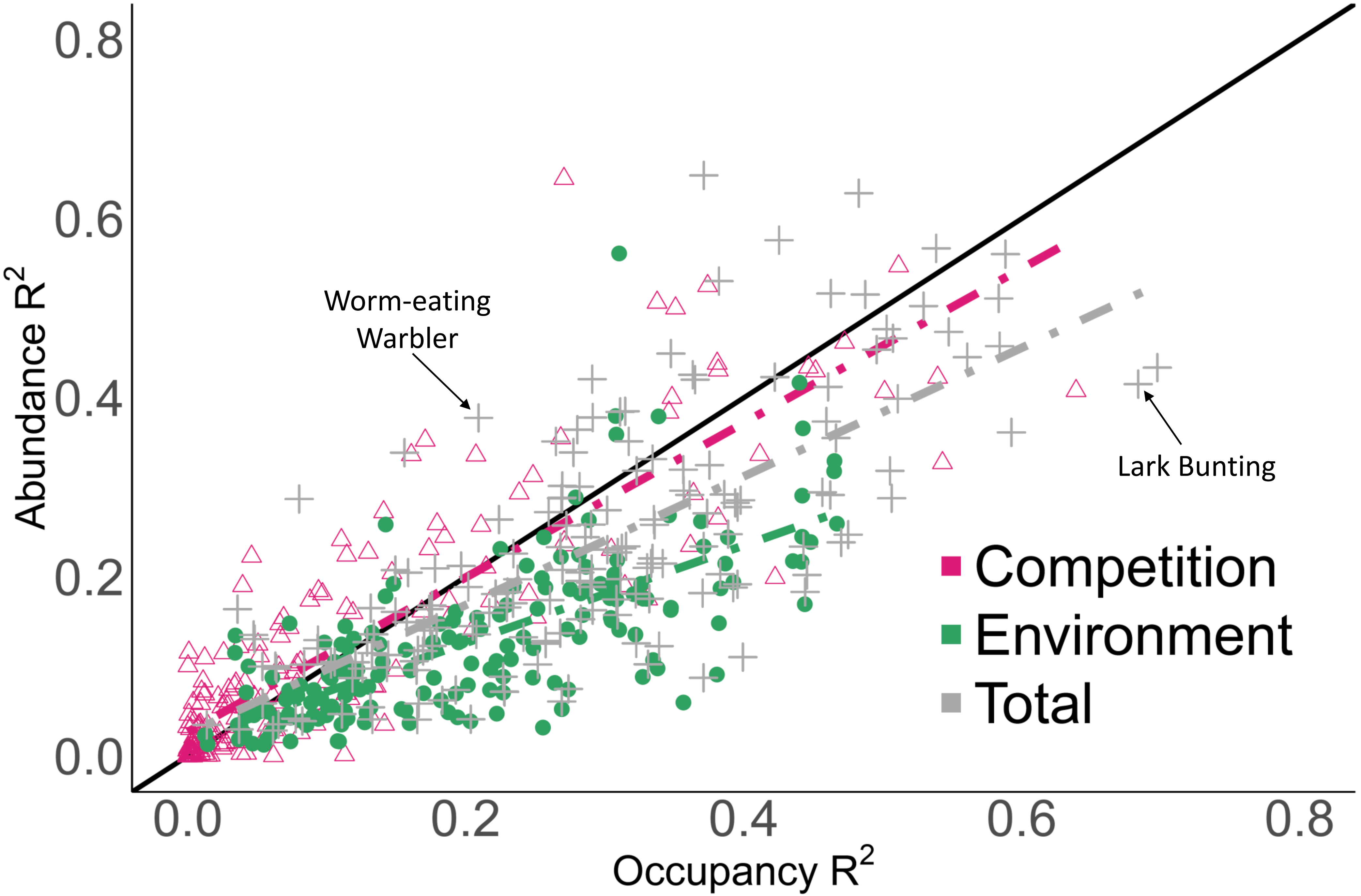
